# Volumetric Denoising: An Approach to Boost Acquisition and Downstream Analysis of Volume Electron Microscopy

**DOI:** 10.1101/2025.08.26.672334

**Authors:** Bohao Chen, Fangfang Wang, Haoyu Wang, Yanchao Zhang, Zhuangzhuang Zhao, Haoran Chen, Hua Han, Xi Chen, Yunfeng Hua

## Abstract

Volume electron microscopy (VEM) enables nanometer-resolution three-dimensional (3D) visualization of biological specimens via serial sectioning and imaging. Owing to precautions against additional challenges in downstream analysis, VEM datasets are often acquired unnecessarily at slow speeds and high resolutions, thereby limiting achievable imaging throughput. By systematically searching for optimal VEM acquisition conditions, we find that sufficient spatial resolution effectively counteracts high image noise in preserving 3D structural information. To further verify that denoising is more effective in restoring volumetric datasets than axial interpolation, machine learning-based methods, including a newly developed 3D context-based denoising model, were compared through various tasks on VEM datasets acquired simultaneously. Our volumetric approach not only outperforms other baseline methods in faithful feature recovery but also facilitates robust serial block-face cutting down to 20 nm by allowing fast imaging. This work provides both an optimized acquisition strategy and volumetric denoising methods as actionable guidelines for maximizing VEM throughput.

## INTRODUCTION

Volume electron microscopy (VEM) is a series of rapidly evolving technologies that enable high-throughput acquisition of volumetric datasets from resin-embedded samples by means of either serial sectioning or iterative block-face ablation.^1–3^ Although a lateral resolution of 5 nm can be readily achieved by all ultramicrotome-based VEM, the minimal cutting thickness that can be sustained over hundreds of consecutive cycles differs from each other due to distinct principles of mechanics.^2,4–6^ Typically, commercial ultramicrotomes produce high-quality serial sections with an interval of 30 or 40 nm using diamond blades, while cutting at smaller steps often leads to increased cutting variability and section damage due to mechanical deformation.^7^ Serial block-face scanning electron microscopy (SBF-SEM), by comparison, enables homogeneous removal of the sample surface at 25 nm intervals with an in-chamber ultramicrotome, because the resin block is simply more rigid than ultrathin sections.^8^ Among all VEM modalities, focused ion beam scanning electron microscopy (FIB-SEM) delivers the greatest axial resolution owing to its ion milling mechanism for precise ablation of the sample surface down to 5 nm.^6,9^ However, given limited setup availability and high imaging costs up-scaled with resolution, the majority of VEM volumes acquired to date – with very few exceptions generated by FIB-SEM – exhibit anisotropic voxel sizes. Although the critical importance of sufficient axial resolution has been previously stressed for both error-free structure annotation (indispensable for the reconstruction of thin neuronal processes) and development of robust automated segmentation algorithms, the impact of spatial resolution on structural information fidelity and automated segmentation performance has not yet been systematically investigated.^4^

As the VEM field pursues ambitious goals of digitizing ever-larger tissue volumes, significant efforts have been devoted to accelerating image acquisition, including the enhancement of sample contrast via saturated heavy metal deposition,^10–12^ the development of high-speed microscopes using camera arrays^13^ or multiple electron beams,^14^ and the establishment of automated, parallelized imaging pipelines.^15^ Nevertheless, when planning practical and cost-efficient imaging workflows on commercial VEM setups, it becomes increasingly important to select optimal (yet empirically determined) acquisition parameters for a good balance between imaging quality and throughput. In addition, improving image signal-to-noise ratio (SNR) via prolonged scanning may significantly compromise achievable axial resolution on SBF-SEM, because reduced cuttability of the resin surface is often attributed to over-exposure to the electron beam.^4,5^ Collectively, this raises a fundamental question facing nearly all VEM acquisitions: should higher priority be given to longer pixel dwell times for improved image SNR, or to smaller pixel sizes and thinner sectioning for higher spatial resolution?

For the above dilemma, recent developments in artificial intelligence (AI)-based image restoration may offer promising solutions. By learning structural features from a variety of VEM datasets, state-of-the-art methods have enabled both image denoising by diffusion^16^ or general^17^ models, and isotropic data generation by nonlinear interpolation.^18–21^ However, the accuracy of model inference has predominantly been evaluated by comparing ground-truth with data recovered from artificial degradations, or by cross-comparisons among different model outputs, leaving the influence of VEM acquisition conditions on model performance largely unvalidated. This motivates us to seek a machine-friendly VEM acquisition strategy for maximizing structural information recovery using the AI model of choice, rather than fine-tuning task-specific models for *post hoc* image restoration.

Here, we used both synthetic and authentic VEM datasets to numerically model the influence of acquisition conditions on structural information fidelity. Next, we compared the performance of several AI models in enhancing the quality of test datasets acquired on a commercial SBF-SEM. We then demonstrated a use-dependent optimization of SBF-SEM acquisition for the most effective image restoration approach. In addition, we validated the resulting improvement in automated image segmentation using datasets enhanced by AI-based methods.

## RESULTS

### Structural information fidelity under different acquisition settings

To quantitatively explore the implications of section thickness and image noise level on VEM dataset quality, we used a publicly available high-quality FIB-SEM dataset of the mouse brain tissue^22^ to generate a series of synthetic datasets through noise masking and spatial down-sampling (Figure 1A). First, the original image stack with an isotropic voxel size of 5 nm was down-sampled in both lateral (XY) and axial (Z) dimensions to yield new stacks at 10-nm and 15-nm voxel sizes. Next, to simulate low-SNR datasets acquired with short pixel dwell times, Poisson-Gaussian noise of appropriate intensity was added to the original images based on the reported reference values^16^ (Figures 1B and S1; also see STAR methods). In parallel, Z-extraction of the original image stack was carried out by sampling every two, three, or four sections, respectively. These generated datasets allow comparisons of dataset quality across combinations of thicker sectioning and faster imaging.

**Figure 1.**
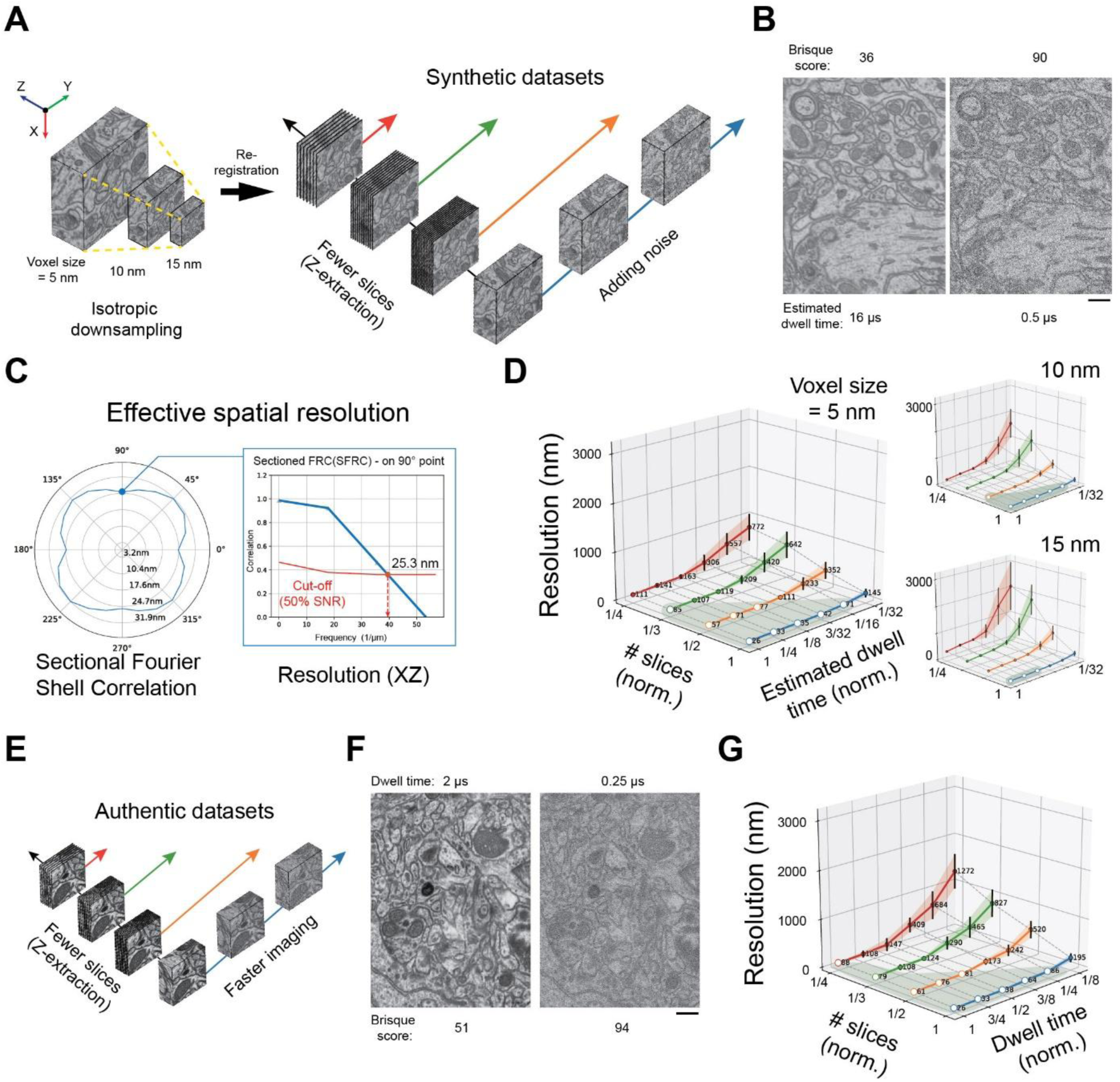
Fidelity of structural information in different VEM samplings: (**A**) Data simulation procedure of a public FIB-SEM volume. Down-sampling yields three synthetic datasets with isotropic voxel sizes. The obtained datasets are progressively masked with noise of varying levels and extracted in the axial direction (Z). **(B)** To the original FIB-SEM images, Poisson-Gaussian noise was added with matched intensity in terms of the BRISQUE (Blind/Referenceless Image Spatial Quality Evaluator) scores reported for specific pixel dwell times as in the reference^16^. Example images masked by different noise levels are shown with corresponding BRISQUE scores, based on which the used dwell times are estimated. Scale bar: 500 nm. (**C**) Quantitative assessment of data quality using the effective spatial resolution, for which a sectional Fourier Shell Correlation (SFSC) is computed for each volume across different angles (as a polar plot). The resolution cut-off in the XZ plane (90° direction) is used to score changes in response to Z-extraction and noise masking. (**D**) Effective XZ resolutions of datasets under different Z-extraction and noise levels. Z-sampling rate (# slices) and noise mask (estimated dwell time) are normalized to the original imaging consumption and scaled approximately to the same fractions. Distinct rising phases of colored lines suggest that resolution worsening in datasets with low slice numbers is significantly accelerated upon heightened noise levels associated with fast scanning. Boundary conditions for the so-called “comfort zone” of acquisition (grey area), which is defined by an effective resolution below 100 nm (open dots). Inset, a similar tendency is observed in modeling with datasets of down-sampled voxel sizes. (**E**) FIB-SEM validation data acquisition procedure of a mouse brain sample with different dwell times and extracted in the axial direction. (**F**) Representative images from the same region acquired at 2.00 µs/pixel (left) and 0.25 µs/pixel (right). The BRISQUE score of the slow-scanned image is 51, and for the fast-scanned one is 94, corresponding to different signal-to-noise ratios. Scale bar: 500 nm. (**G**) Effective XZ resolutions of validation datasets under different Z-extraction and imaging dwell times.

Quality assessments of these synthetic datasets were performed by computing their effective spatial resolution (Figures 1C and S2). As previously demonstrated^23,24^, this metric provided a way to objectively compare volumetric data quality by measuring the normalized cross-correlation within individual Fourier shells (see STAR Methods). As the VEM acquisition duration of a given volume scales approximately with axial step size and scanning speed, we normalized the section numbers and the estimated acquisition times to facilitate a comparison across acquisition conditions with combined parameters (Figure 1D). This revealed that the effective XZ resolutions obtained from data with a high Z-sampling rate were robust to varying image SNRs, forming a “comfort zone” for relatively stable data quality under different scanning speeds. Consistently, it was observed across all simulated scenarios of further data down-sampling (Figure 1D inset). To rule out artificial image degradation-induced effects, authentic datasets were acquired from a brain sample at different scanning speeds using plasma-FIB-SEM (Figure 1E-1G). On these datasets, we validated the differential contributions of axial sampling rate and image SNR to the effective resolution of a volumetric dataset.

To answer whether the degradation of VEM data quality may affect the organelle segmentation, the accuracy of automatic mitochondrial segmentation was compared across the synthetic FIB-SEM datasets (Figure 2A). We found that even with a modest image SNR, low segmentation errors (1-IoU) were achievable from datasets of a high Z-sampling rate (Figure 2B), arguing for the importance of spatial sampling rate to automatic organelle segmentation. Furthermore, we show that the improved effective resolution led to a decline in segmentation error (Figure 2C).

**Figure 2.**
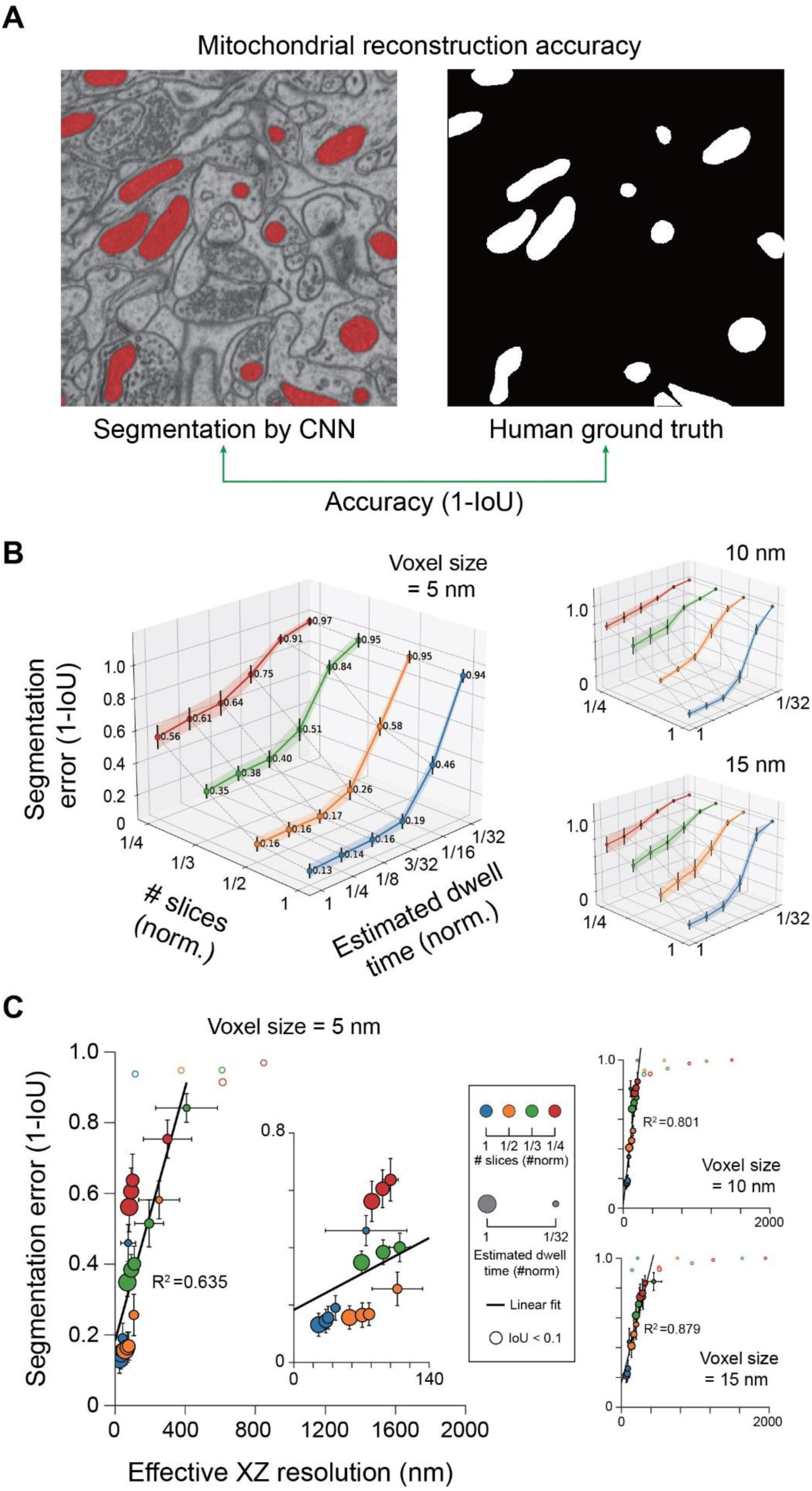
VEM data quality influences the automated organelle segmentation: (**A**) Data quality assessment by segmentation accuracy of mitochondria using the convolutional neural network (CNN)-based model for mitochondria. The segmentation accuracy is scored using the Intersection-over-Union (IoU) between model inference and human annotation. **(B)** CNN segmentation error (as 1 – IoU) on synthetic datasets reveals declined model performance upon reduced Z-sampling and shortened dwell time. Inset: Similar scenarios for datasets with down-sampled voxel sizes. **(C)** Plot of CNN segmentation error (1-IoU) of mitochondria against the data effective resolution (data from Fig. 1). Linear fit, R² > 0.635 (outliers: IoU < 0.1). Inset: same plot with the datasets of 10 nm (top) and 15 nm (below) voxel size. These results demonstrate a strong positive correlation between the data’s effective resolution and subsequent 3D reconstruction quality, validating the usage of the former as a general metric for data quality assessment.

### Machine learning-based restoration of serial images

The above results substantiate an underestimated role of section thickness in maintaining VEM data quality by counteracting image noise, but it remains unclear whether the axial interpolation of image stacks will be more challenging for computer algorithms than denoising. To test this, we designed a hybrid imaging routine to acquire interlaced reference (high-SNR) and snapshot (low-SNR) slices by alternating scanning speeds. Specifically, a mouse brain sample was cut consecutively at a nominal thickness of 50 nm and imaged redundantly at two different pixel dwell times (2 µs and 0.5 µs) using a commercial SBF-SEM (Figure 3A). The acquired images were arranged to generate distinct test datasets (Figure 3B), including a reference stack (R-stack, scanned at 2 µs/pixel), a snapshot stack (S-stack, scanned at 0.5 µs/pixel), a subset of reference stack (½R-stack, skipping every other slices), and a hybrid dataset (S/r-stack, a fixed combination of ¾ snapshot and ¼ reference slices). These datasets provided a unique opportunity to compare the model performance for interpolation and denoising based on a shared ground truth (Figure 3C).

**Figure 3.**
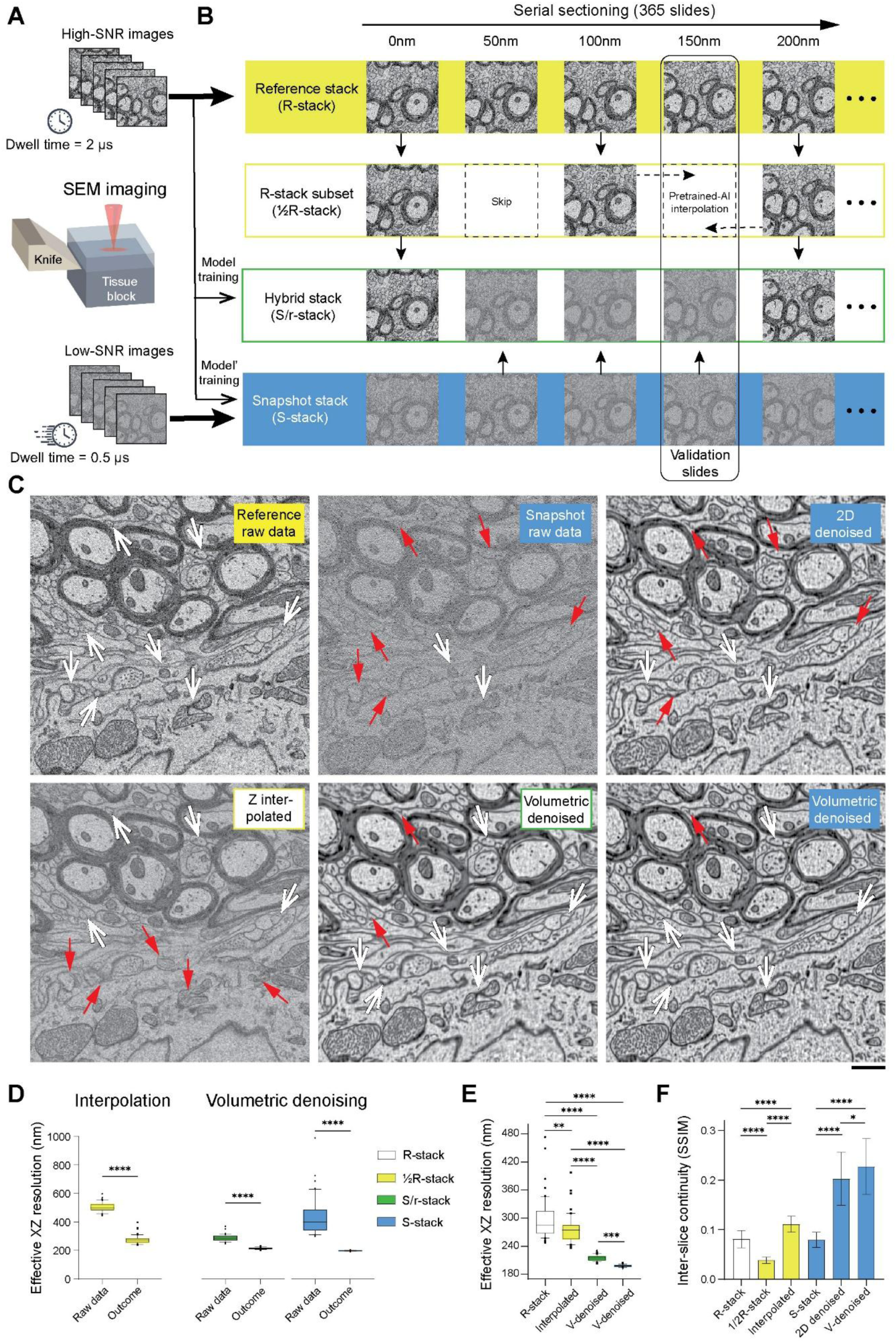
Denoising of image stack using 3D structural context: (**A**) Schematic illustration of a hybrid image acquisition using SBF-SEM. Repeat block-face scanning produces image pairs of low and high signal-to-noise-ratio (SNR). (**B**) Full stack of high-SNR images (R-stack, yellow solid) serves as a reference, whereas the snapshot dataset containing exclusively low-SNR images (S-stack, blue solid) is for the method development. Two synthetic image stacks: For the ½R-stack (yellow box), every other slow-scan-only slice is replaced with an interpolated virtual slice. S/r-stack (green box) is made up of fixed combinations of one reference and three snapshot slices. (**C**) Representative image of the same slice in the reference stack (up left) and snapshot dataset (up middle), as well as AI-denoised S-stack only using 2D information (up right). Similarly, the same slice was generated by 3D methods, including AI-based interpolation (bottom left) as well as supervised volumetric (v-) denoising on the S/r-stack (bottom middle) and the S-stack (bottom right). Arrows indicate faithfully preserved structures (white) and artifacts (red). Scale bar, 1000 nm. (**D**) Improvement of effective resolution by AI-based image restoration. (Left) By interpolation, the effective resolution for the ½R-stack drops from 502.3 ± 32.7 nm to 275.9 ± 31.2 nm. The v-denoising improves the effective resolution for the S/r-stack (middle) from 287.2 ± 23.4 nm to 214.8 ± 6.5 nm, as well as for the S-stack (right) from 433.1 ± 132.6 nm to 197.9 ± 2.6 nm. (**E**) Comparison of the achieved effective resolutions by different 3D approaches from (D) and ground truth (R-stack). Effective resolution of R-stack: 297.1 ± 45.3 nm. **(F)** Quantification of inter-slice structural continuity in datasets containing raw, interpolated, and denoised images. AI-interpolated ½R-stack achieved a higher score in Structural Similarity Index Measure (SSIM) than both the ½R-stack and the ground truth (R-stack). Compared to raw data (S-stack), both 2D and v-denoised datasets demonstrated significantly improved structural continuity, with the highest SSIM observed in the v-denoised dataset.

Moreover, since most cellular structures appear continuous across consecutive ultrathin sections, we speculate that axial oversampling may provide mutual verification of true signals based on 3D structural priors that can be used by AI models for volumetric data restoration. Thus, we developed a volumetric denoising model (Figure S3, see STAR Methods for more details) and added it to the method comparison. First, apparently fewer artifacts and blurred regions were observed in the denoised outputs of the S/r– and S-stacks when leveraging additional 3D structural information. Second, the incorporation of high-SNR reference frames in the input (S/r-stack versus S-stack) seemed not to improve outputs of the volumetric denoising model. Third, we observed better structural preservation – with less aliasing and blurring – in the denoised images (S/r– and S-stack) than in the interpolated ½R-stack, again underscoring the pivotal role of sufficient axial sampling beyond image SNR. These observations were confirmed quantitatively by comparing the effective resolution across different model outputs (Figure 3D). Surprisingly, volumetric denoising yielded a slightly lower data quality for S/r-stack than for S-stack (Figure 3E). As a plausible explanation for that, the volumetric denoised S/r stack retained the original reference frames, which had a higher noise level than those in the volumetric denoised S-stack (Figure S4). Most importantly, the inter-slice structural continuity was maximally preserved in the volumetric denoised image series compared to interpolated or 2D (frame-based) denoised outputs (Figure 3F).

### Optimization of VEM acquisition for volumetric denoising

As the axial resolution of SBF-SEM is mainly limited by mechanical cutting of the resin block, imaging with a short pixel dwell time may be favorable for cutting at small step sizes (Video S1). In turn, the model performance of volumetric denoising may benefit from an increased axial sampling rate. To test this concept, we acquired three image stacks from the same sample using a 0.5 µs dwell time but at different nominal cutting thicknesses (50 nm, 25 nm, and 20 nm). Instead of using high-SNR images as references, we adopted a self-supervised strategy for model training. Although the AI model delivered visually comparable results for image stacks acquired at all tested cutting thicknesses (Figure 4A-4C, Videos S2-S3), quantification revealed differential improvements in the effective XZ resolution from 850.3 ± 361.8 nm to 209.5 ± 6.6 nm (50-nm cutting), 306.5 ± 98.9 nm to 104.1 ± 3.3 nm (25-nm cutting), and 263.6 ± 97.0 nm to 83.6 ± 2.0 nm (20-nm cutting) by self-supervised volumetric denoising (Figure 4D). Note that the effective XZ resolution obtained after denoising is linearly correlated with the nominal cutting thickness (Figure 4E), serving as a good indication of minimal residual noise in the model outputs. We also controlled inter-sample variability by comparing the XY component of effective resolution, which is insensitive to cutting thickness (Figure S5). Note that our volumetric denoising minimized the occurrences of unrealistic, sudden structural changes between consecutive slices in comparison to the outcomes of frame-wise approaches (Figure 4F). However, our volumetric denoising yielded a significant increase in inter-slice structural continuity for datasets up to a cutting thickness of 75 nm, beyond which the advantage of using 3D structural information was subtle in denoising VEM datasets compared to frame-based 2D methods (Figure 4G).

**Figure 4.**
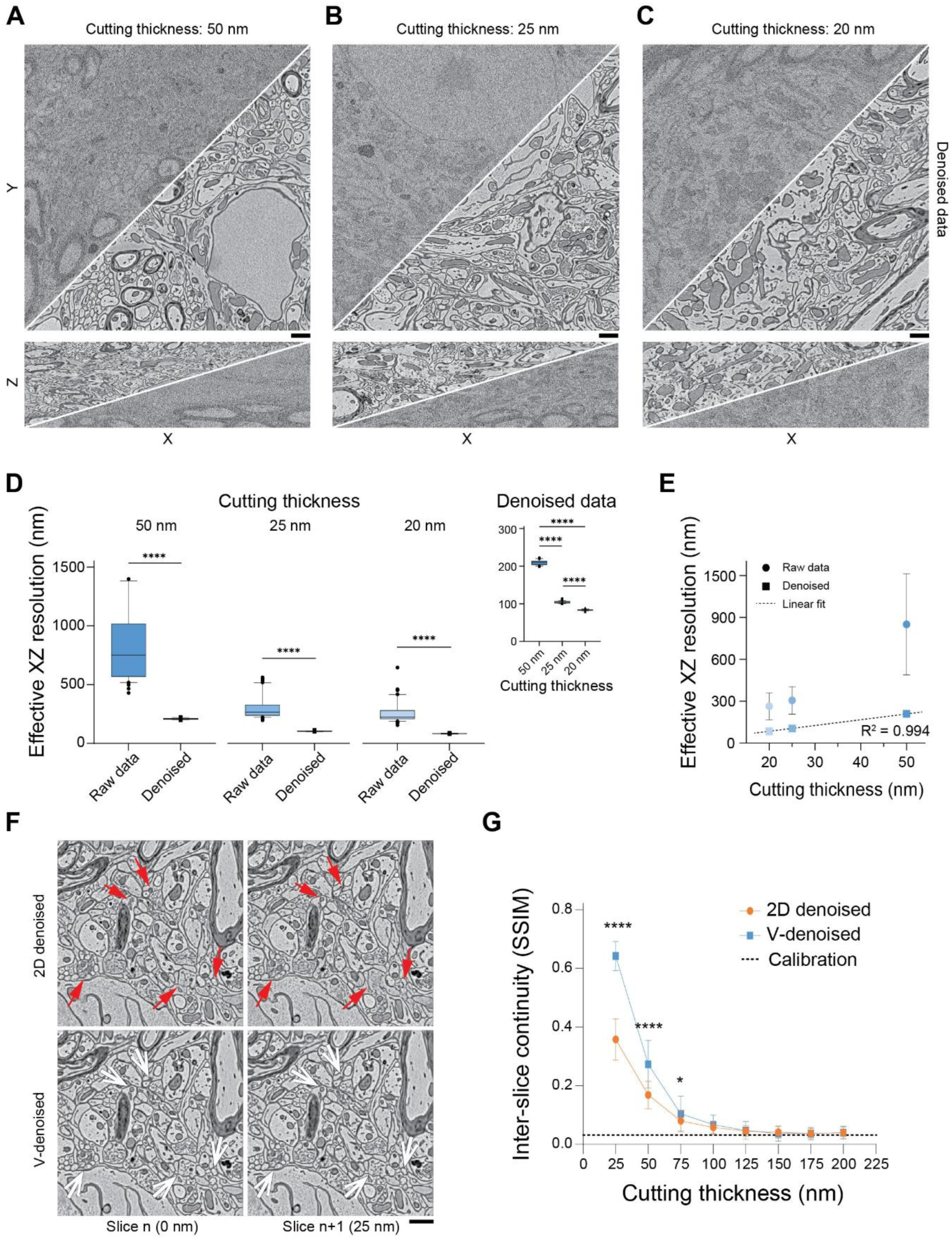
Improved Z-sampling allowed by volumetric denoising: (**A-C**) Three SBF-SEM datasets acquired in the snapshot mode (0.5 µs/pixel) but at different cutting thicknesses (A: 50 nm, B: 25 nm, C: 20 nm) for the same sample. Representative raw image and volumetric denoised image by a self-supervised model are shown in both XY (top) and XZ (bottom) views. Scale bar, 1 µm. (**D**) Differential improvements in the effective XZ resolution after denoising are observed for datasets of different cutting thicknesses (50-nm: from 850.3 ± 361.8 nm to 209.5 ± 6.6 nm; 25nm: from 306.5 ± 98.9 nm to 104.1 ± 3.3 nm; 20-nm: from 263.6 ± 97.0 nm to 83.6 ± 2.0 nm). Inset: Plot of the achieved XZ resolution from d. against physical cutting thickness. (**F**) 2D denoising (top row) and v-denoising (bottom row) were contact on the same dataset, and representative results of two consecutive slices (Slice n and Slice n+1) were shown. Upon a 25 nm cut, independent 2D denoising leads to more frequent sudden changes (red arrows), which are considered irrational and not observed in the v-denoised dataset (white arrows). Scale bar, 1 µm. (**G**) Measures of inter-slice structural continuity using SSIM as a function of cutting thickness. Datasets of increasing thicknesses (25 nm to 200 nm) were generated from the 25-nm dataset. Results of 2D denoising (blue circles) and v-denoising (orange squares) were compared. The advantage of v-denoising over 2D denoising in terms of preserving continuous structures is only observed in datasets with a slice thickness of less than 75 nm. Calibration of random structure continuity was carried out between two completely different patches within the volume (dotted line).

### Volumetric denoising facilitates downstream VEM analysis

To find out whether volumetric denoising contributes to downstream analysis tasks, such as organelle segmentation and large-scale reconstruction, we used a plasma-FIB-SEM to acquire two volumetric datasets from the same field of view on a mouse brain sample at different scanning speeds (0.5 and 2.0 µs/pixel). To obtain the ground-truth, mitochondria and synapses were manually annotated in the slow-scanned reference images. Pretrained AI models for mitochondria and synapse segmentation were applied to the raw snapshot dataset acquired and the corresponding denoised outputs from four different methods, including the local smoothing in FIJI^25^, 2D self-supervised denoising (Blind2Unblind)^26^, self-supervised movie denoising (DeepSeMi)^27^, and ours (v-denoising, Figure 5A). By comparing to the ground-truth annotation, both mitochondria and synapse segmentation models achieved higher accuracy on the dataset denoised by the volumetric approach than the other tested baseline methods (Figure 5B and 5C). These results demonstrate that volumetric denoising can maximally improve the performance of existing segmentation models by providing accurately restored fine structural features.

**Figure 5.**
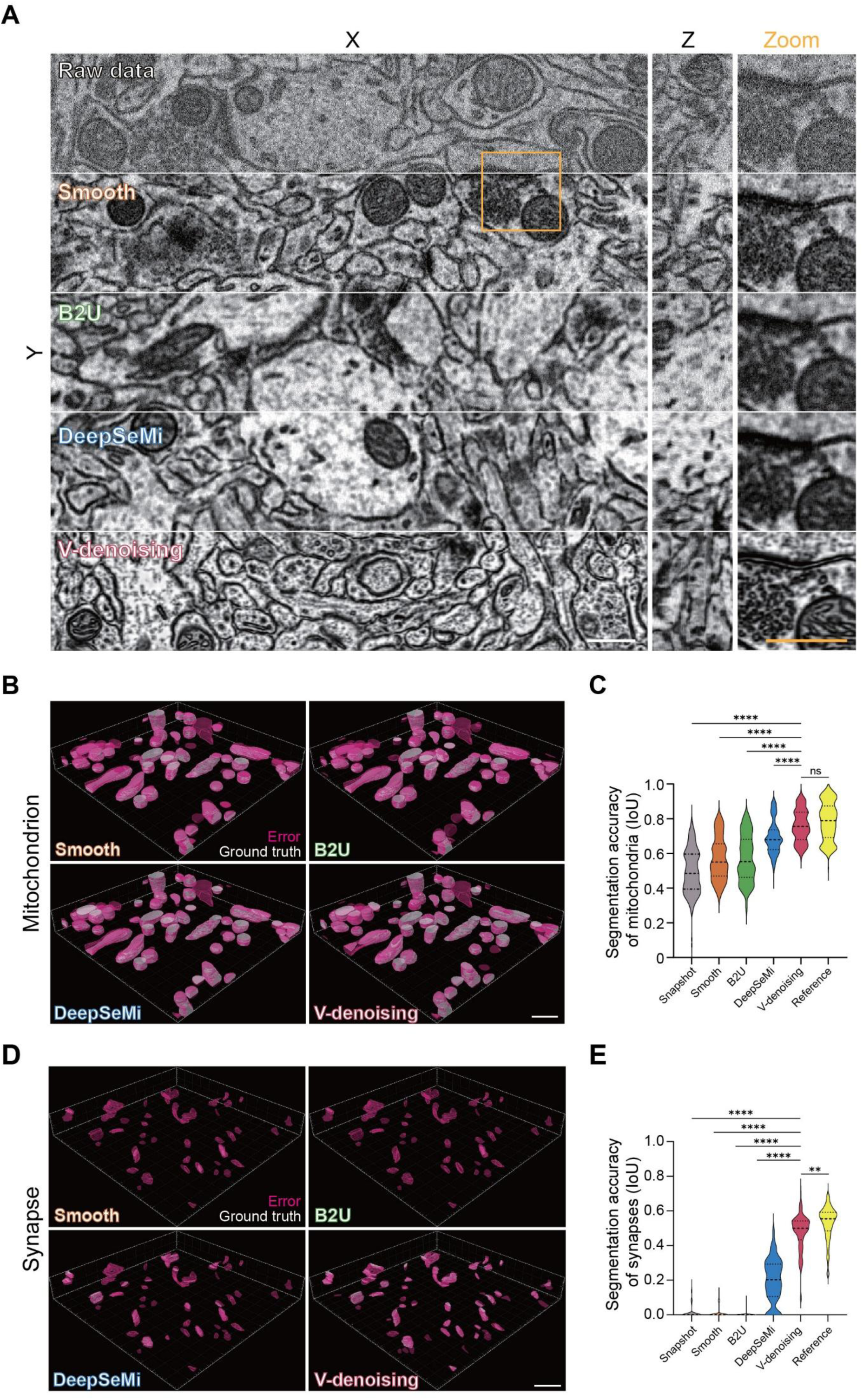
Volumetric denoising improves segmentation accuracy of organelles: **(A)** Denoising the plasma FIB-SEM dataset of a mouse brain sample using different methods. Volumetric denoising (v-denoised), 3-by-3 smoothing (smooth), Blind2Unblind (B2U), and DeepSeMi were applied on the fast-acquired dataset (0.5 µs/pixel and 5 nm isotropic voxel size). Inset: enlarged region of interest as indicated (yellow box). Scale bar, 500 nm. **(B)** The same mitochondrion instances were obtained from automatic segmentation on differently denoised datasets. Ground truth (white) and test segmentations (magenta) are spatially overlaid. Scale bar, 1 µm. **(C)** Segmentation accuracy of mitochondria on the raw dataset (snapshot), denoised datasets, and the reference dataset. Model outputs are compared with manual annotation on the reference dataset as ground truth. Intersection over union (IoU): 0.496 ± 0.130 (snapshot), 0.570 ± 0.123 (smooth), 0.571 ± 0.134 (B2U), 0.692 ± 0.096 (DeepSeMi), and 0.759 ± 0.091 (v-denoised), 0.785 ± 0.110 (reference). Friedman test: p < 0.0001 (snapshot vs. v-denoised), p < 0.0001 (smooth vs. v-denoised), p < 0.0001 (B2U vs. v-denoised), p < 0.0001 (DeepSeMi vs. v-denoised), and p = 0.299 (v-denoised vs. reference). **(D)** Example of segmented synapse from the same datasets as in B. Scale bar, 1 µm. **(E)** Segmentation accuracy of synapses on the raw and denoised datasets. IoU: 0.019 ± 0.043 (snapshot), 0.010 ± 0.028 (smooth), 0.002 ± 0.014 (B2U), 0.197 ± 0.125 (DeepSeMi), and 0.477 ± 0.108 (v-denoised), 0.529 ± 0.100 (reference). Friedman test: p < 0.0001 (snapshot vs. v-denoised), p < 0.0001 (smooth vs. v-denoised), p < 0.0001 (B2U vs. v-denoised), p < 0.0001 (DeepSeMi vs. v-denoised), p = 0.0082 (v-denoised vs. reference).

In addition, we applied the volumetric denoising to a sub-volume (1000 × 1000 × 1000 voxels, 12 × 12 × 35 nm^3^) from our previously published SBF-SEM dataset of the zebrafish spinal cord^28^ and ran an automated neuronal segmentation pipeline separately on the raw and denoised datasets (Figure 6A). For quantification, seven neurites were randomly selected in both image stacks for manual proofreading, and the total annotation time required by an expert annotator was compared (Figure 6B). We found that the denoising significantly reduced the occurrence of neurite over-segmentations (Fig. 6C, raw data: 5.607 ± 6.530 / µm, denoised data: 3.950 ± 5.264 / µm), thereby effectively saving 29% human work (reduced from 1.01 to 0.72 min/µm) involved in merging segmentation and error correction (Figure 6D). Given that manual proofreading has become the highest cost and major bottleneck in large-scale connectomic studies, integrating the volumetric denoising into the data analysis workflow prior to segmentation may yield substantial benefits by reducing human labor requirements. Together, these results demonstrate the broad applicability of our self-supervised volumetric denoising model across authentic datasets from different experimental setups and VEM modalities, robustly facilitating a wide range of downstream analysis tasks.

**Figure 6.**
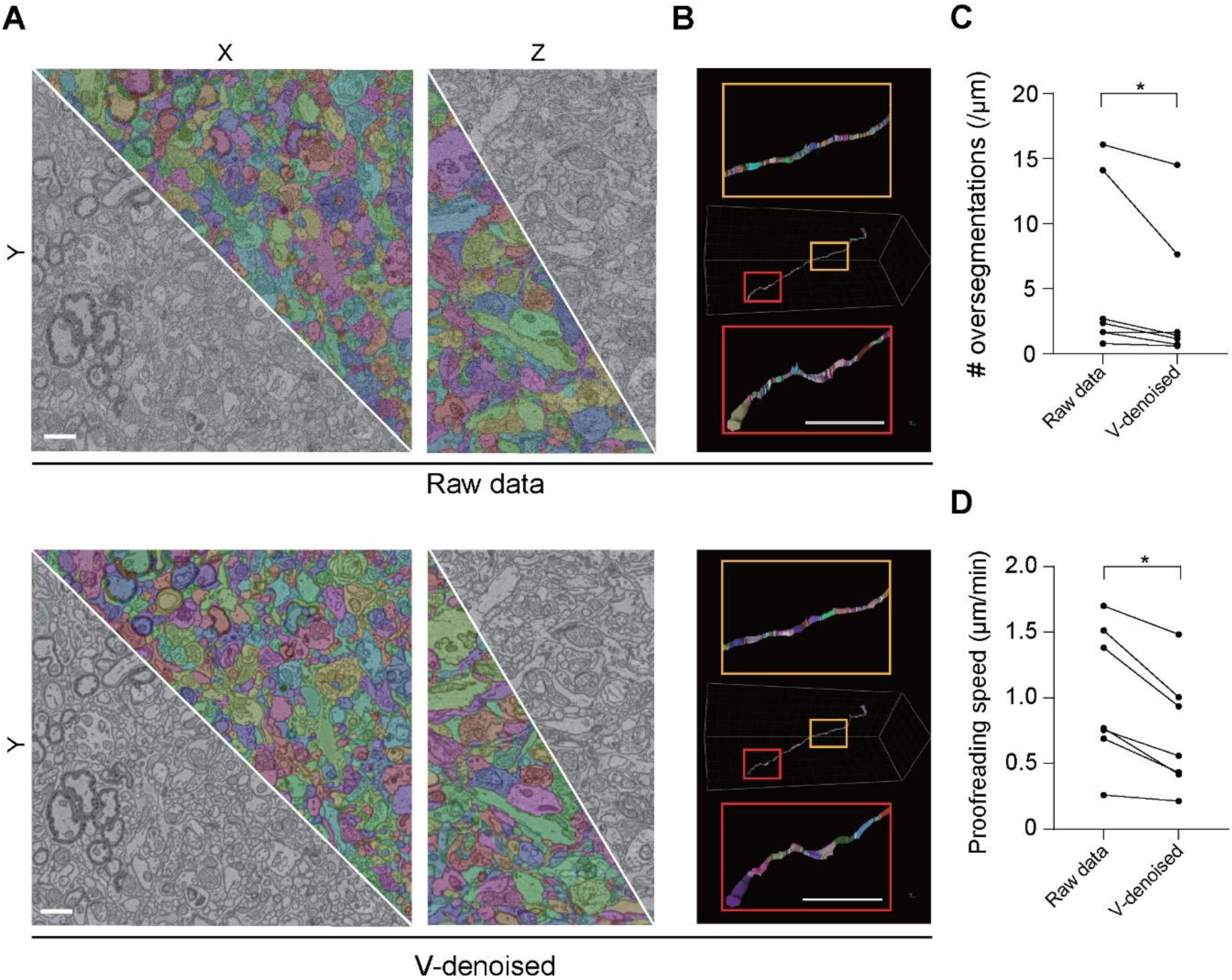
Volumetric denoising accelerates neurite reconstruction: **(A)** Automatic segmentation and neurite reconstruction test on the raw (top) and volumetric denoised datasets (below) of the zebrafish spinal cord. Scale bar: 1 µm, **(B)** An automatic segmented neurite example with over-segmentations (pseudo-colored) in the raw (top) and denoised datasets (below) before proofreading. Scale bar: 1 µm. **(C)** The counts of neurite oversegmentation (normalized to pathlength) in the raw and denoised datasets. Raw data: 5.607 ± 6.530 / µm, v-denoised: 3.950 ± 5.264/µm, n = 7, paired Wilcoxon test: p = 0.016. **(D)** The normalized proofreading time required for individual test neurites. Raw data: 1.011 ± 0.525 min/µm, v-denoised 0.722 ± 0.441 min / µm, n = 7, paired Wilcoxon test: p = 0.031.

## DISCUSSION

In principle, VEM imaging of biological specimens is a 3D-digitalization process, in which spatial and temporal sampling rates correspond to voxel size and dwell time, respectively. Unlike light microscopy, the lateral resolution of modern electron microscopy is typically more than sufficient for visualizing most cellular structures, producing oversampled structural details in electron micrographs. By contrast, technical challenges or time constraints in sample sectioning at smaller step sizes lead to an under-sampling in the axial direction, strongly influencing the VEM data quality. By quantifying the changes in VEM dataset quality upon trade-offs between axial sampling rate and pixel dwell time (Figure 1D and 1G), we provide a fact-based argument for the pivotal role of sufficient spatial sampling in maintaining structural information fidelity over high image contrast. Besides, the boundary of combined acquisition parameters defines a so-called “comfort zone”, within which the sampling-dependent improvement in data quality reaches a plateau. Although tissue-specific calibration may be required, our finding is particularly instructive for the selection of imaging parameters when this trade-off must be made. Notably, emerging techniques, such as gas-cluster ion beam^29^ and parallel ion beam source,^30^ are capable of reducing the thickness of large-scale surface etching to 10-nm precision, paving the way for industrial-scale tissue mapping at desired sampling rates in the axial direction.

Moreover, our quantitative comparison of AI performance across various image restoration tasks suggests that volumetric denoising outperforms interpolation in real experiment settings (Figure 3D). This finding is significant in two aspects. First, it provides proof-of-principle evidence that AI can learn spatially continuous structural features to accurately distinguish stable signals from random noise in data volumes. Second, as computational systems are far more powerful than humans at memorizing and comparing large image sequences over a wide contrast range, this stimulates new thinking of VEM sampling strategies that should be tailored for AI-based image processing rather than human visual assessment. Indeed, visually favorable high-SNR reference images are found to be not essential for accurate volumetric denoising by the trained AI model (Figure 3E). Despite its redundant role in improving the model performance, our hybrid acquisition approach may serve as a human-in-the-loop paradigm for the future development of AI-based image processing methods beyond denoising, for instance, super-resolution or defocus correction.

As self-supervised volumetric denoising no longer requires the reference frames (Figure 4), we expect broad applications on VEM volumes of different tissue types and experimental setups with minimal data-specific model tuning. More importantly, it provides a new perspective for serial block-face imaging to improve axial resolution and surface removal quality. In the practice of both SBF and FIB-SEM, electron beam-induced radiation damage occurs when imaging with prolonged pixel dwell times or even focusing with repeated scans. As resin in the interaction volume degrades, the sample surface is progressively lowered, therefore hindering the subsequent surface removal. To the best of our knowledge, the cutting thickness of SBF-SEM is currently limited to 25 nm,^31–33^ while our AI-denoising enables data acquisition with snapshots and reliable cuts at a step size of 20 nm using a commercial setup (Figure 4E). With reduced intersection differences owing to smaller cutting thickness, we anticipate an improved Z-alignment of the image stack, which may further contribute to the performance of volumetric denoising.

Lastly, this study provides an example of using AI-based image restoration to optimize the data acquisition of VEM, rather than a *post hoc* image enhancement application. Note that the hybrid acquisition routine proposed in this study allows an arbitrary combination of snapshot and reference frames, which is fundamentally different from conventional fixed-parameter VEM acquisition schemes. This flexible imaging mode has the potential to automatically adapt application-specific trade-offs between throughput and image quality while maintaining biologically meaningful resolution. Most recently, a specific AI model has been developed to predict regions containing synapses and thin neurites for subsequent high-resolution local rescan with prolonged dwell time.^34^ In a similar fashion, future work may integrate the volumetric denoising model into an SBF-SEM or FIB-SEM system, so that imaging parameters can be updated based on model outputs in real time. Ultimately, this will accelerate the VEM data collection with minimal human involvement.

In summary, the present work demonstrates through quantitative analysis that sufficient VEM spatial sampling plays a pivotal role in effectively preserving structural information. Furthermore, we deliver a machine learning-based volumetric denoising approach for image stack restoration with superior accuracy. Cross-dataset and cross-VEM modality validations confirm that the volumetric denoising method benefits not only the axial resolution improvement of SBF-SEM but also a variety of VEM downstream analysis tasks. Furthermore, the concept of volumetric denoising may be applicable across other high-throughput imaging modalities for connectomics, including X-ray tomography^35^ and possibly expansion light microscopy, like LICONN^36^.

### Limitations of the study

This study demonstrates the concept of using 3D structural context to achieve accurate denoising of image stacks. Although the model performance depends primarily on the spatial sampling rate, the maximum noise level for an error-free image restoration remains to be further characterized. Thus, we strongly recommend monitoring the model’s performance during VEM acquisition, for example, by incorporating checkpoints to routinely acquire high-SNR reference images for validation of denoising accuracy.

## RESOURCE AVAILABILITY

### Lead contact

Further information and requests for resources and reagents should be directed to and will be fulfilled by the lead contact, Yunfeng Hua.

### Material availability

This study did not generate new unique reagents.

### Data and code availability

- Raw VEM datasets and segmentations supporting the current study are publicly available via the links listed in the key resource table.
- The source codes used in this paper are available as of the publication date ( https://github.com/hmzawz2/OptimizedAcquisition4vEM, <u>d8cb2f6</u>).
- Any additional information required to reanalyze the data reported in this paper is available from the lead contact upon request.

## ACKNOWLEDGEMENTS

We thank Prof. Ming Lei and Prof. Guoqiang Bi for the discussion. This study was supported by the Scientific Research Instrument and Equipment Development Project of Chinese Academy of Sciences (PTYQ2025TD0002 to H.H.), the National Natural Science Foundation of China (32171461 to H.H., 82171133 to Y.H.), and the Innovative Research Team of High-level Local Universities in Shanghai (SHSMU-ZLCX20211700).

## AUTHOR CONTRIBUTIONS

Y.H., X.C. conceived the study; B.C. conducted the study under the supervision of Y.H., X.C., H.H.; The software was developed by B.C with the help of Y.Z., H.C., ZZ; B.C., H.W., H.C., Z.Z. performed the analysis; F.W. performed the SBF-SEM experiment; Y.H., H.H. acquired funding; Y.H., B.C. drafted the manuscript with contributions from all authors.

## STAR METHODS

### EXPERIMENTAL MODEL AND STUDY PARTICIPANT DETAILS

Adult C57BL/6 mice (male) were purchased from Shanghai Jihui Experimental Animal Feeding Co., Ltd. (Shanghai, China). All procedures were approved by the ethical committee of Shanghai Ninth People’s Hospital (No. SH9H-2019-A567-1) and conducted at Shanghai Institute of Precision Medicine.

## METHOD DETAILS

### Sample preparation

The mouse brain samples were prepared according to the previously published procedure^10^ with minor modifications. In brief, animals were anesthetized with isoflurane inhalation and transcardially perfused with 30 mL 1.5 M cacodylate buffer (CB, pH 7.4), followed by an ice-cold fixative mixture containing 2% paraformaldehyde and 2.5% glutaraldehyde in 0.08 M CB. The dissected brain tissues were post-fixed at 4℃ in the same fixative for 48 hours and then sliced 500-µm thick using a vibratome (Leica VT 1200S) in 0.15 M CB. The *en bloc* staining was done by multiple rounds of incubation with heavy metals, including 2% OsO_4_ (in 0.15 M CB), 2.5% ferrocyanide (in 0.15 M CB), 2% OsO_4_ (in 0.15 M CB) at room temperature (RT) for 1.5, 1.5, and 1.0 hours, respectively. After being washed with 0.15 M CB and water for 30 min each, the samples were incubated at RT in thiocarbonhydrazide (saturated aqueous solution) for 1 hour at RT, 2% OsO_4_ aqueous solution for 1.5 hours at RT, and 1% uranyl acetate (aqueous solution) for 2 hours at 50℃ with an intermediate washing step (twice in water for 30 min each). The sample dehydration was carried out through a graded acetone series (50%, 75%, 90%, 30 min each, all cooled at 4℃) and in pure acetone (100%, three times, 30 min each). Finally, the samples were infiltrated through a 1:1 mixture of acetone and Spurr’s resin monomer (4.1g ERL 4221, 0.95g DER 736, 5.9g NSA, and 1% DMAE) overnight on a rotator, then embedded in pure resin for 8 hours before being baked in a 70℃ oven for 72 hours.

### Data acquisition of SBF-SEM

The sample was mounted on an aluminum metal rivet using epoxy adhesive for 12 hours and trimmed to a rectangular shape (0.8 × 0.8 mm^2^) using a sample trimmer (Leica, TRIM2). Next, the sample surface was prepared by a diamond knife using an ultramicrotome (Leica, UC7). Sputter coating of a thin layer of gold (Leica, EM ACE600) was applied to the sample block, which was then imaged using a field-emission SEM (Zeiss, Gemini300) equipped with an in-chamber ultramicrotome (Gatan, 3View2XP) and back-scattered electron detector (Gatan, OnPoint), with a beam current of 177 pA measured on the instrument.

The SEM imaging was carried out with an incident beam energy of 2 keV and a pixel size of 15 nm. For Fig. 3, serial images (4000 × 4000 pixels, 365 slices) were acquired at a dwell time of 0.5 μs with a cutting thickness of 50 nm. For each imaging and cutting cycle, repeated scans with a dwell time of 2.0 μs were manually triggered to capture redundant images of improved quality. These serial images were assigned in use-dependent orders to compose the reference stack (all frames acquired with 2.0-μs dwell time), the snapshot stack (all frames acquired with 0.5-μs dwell time), and the fast-slow-scan dataset (a combination of one 2.0-μs-dwell-time frame and three 0.5-μs-dwell-time frames). For Fig. 4, the snapshot stack was reused for the 50-nm-cutting experiment. And two independent datasets were acquired using the same imaging parameters, except that the cutting thickness was reduced to 25 nm (902 slices) and 20 nm (626 slices). All SBF-SEM datasets contained at least 301 slices for model training and testing after Z-alignment using a self-written MATLAB script based on cross-correlation maximum between consecutive slices before analysis.^37^

### Data acquisition of plasma-FIB-SEM

The authentic datasets of the same mouse cortex sample under different acquisition conditions were generated using a plasma-FIB-SEM (Thermo Scientific Hydra Bio). Incident beam energy of the SEM was 2 keV, and a back-scattered electron detector was used for signal collection. For surface ablation, an oxygen plasma source with a milling current of 45 nA was used.

In Figure 1E, six datasets (332 sections of 2048 × 2048 pixels) were acquired at an isotropic voxel size of 5 × 5 × 5 nm³ from the same field of view. Each block-face was scanned repeatedly at dwell times of 0.25, 0.5, 0.75, 1.0, 1.5, and 2.0 µs, respectively. To generate datasets with reduced axial sampling, sections were excluded with required intervals. In Figure 5A, two overlapped datasets (181 sections of 1300 × 1300 pixels) were acquired from the same sample with 0.5 µs (snapshot) and 2.0 µs (reference) dwell times at a voxel size of 5 × 5 × 5 nm³.

### Data simulation on a public FIB-SEM volume

To simulate dataset quality under different image settings (Figure 1), a publicly available dataset was used, which was acquired from the mouse CA1 hippocampus region using FIB-SEM at an isotropic voxel size of 5 nm (https://www.epfl.ch/labs/cvlab/data/data-em/).^22^ For the XY down-sampling, binning of image pixels at a factor of 2 and 3 was used, yielding a final XY pixel size of 10 and 15 nm. To perform down-sampling along the Z-direction at a factor of k, the new image stack kept every k slice from the original stack. For a given volume, this process resulted in a k-fold increase in the slice thickness, a k-fold reduction in the number of imaged slices, and an equivalent 1/k fraction of the original acquisition time.

To simulate the effect of a reduced pixel dwell time, we manipulated the image signal-to-noise ratio by introducing Poisson-Gaussian noise of different levels to the original images. The outcomes were scored using the Blind/Referenceless Image Spatial Quality Evaluator (BRISQUE)^38^, implemented via the pyiqa (https://github.com/chaofengc/IQA-PyTorch<u>, v0.1.14.1</u>) for a quantitative calibration. In detail, the variance of the noise used for masking was adjusted so that the obtained BRISQUE score of the masked image matched closely the documented value for images acquired with a specific dwell time.^16^ By doing so, we were able to generate a series of datasets with comparable noise levels to images scanned with different dwell times of 0.5 μs, 1.0 μs, 1.5 μs, 2.0 μs, 4.0 μs, and 16.0 μs, respectively (Fig. S1).

### Computing effective spatial resolution of volumetric datasets

Sectional Fourier Shell Correlation (SFSC) analysis was used to estimate the effective spatial resolution of the cubed VEM stacks.^24^ This method determined directional resolution by calculating the normalized cross-correlation within wedge-shaped sectors of individual Fourier shells. For analysis of an EM volume, the input data was first split into two pairs of sub-volumes based on pixel coordinate on the XY plane:

Pair 1: One sub-volume containing all pixels at (even, even) coordinates and the other containing pixels at (odd, odd) coordinates.

Pair 2: A second pair formed from pixels at (even, odd) and (odd, even) coordinates.

The SFSC for each pair was then calculated using the normalized cross-correlation between their Fourier transforms:

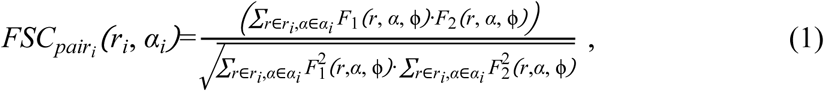

where *F*_1_(*r*, *α*, ϕ) and *F*_2_(*r*, *α*, ϕ) denote the voxels in two Fourier-transformed sub-volumes within pair*_i_* that are located at a given distance *r_i_* from the origin and within an orientation sector, defined by *α* and ϕ. Finally, the SFSC curves on each pair were averaged to yield the final SFSC measurement for the input volume:

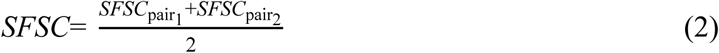

To ensure computational efficiency while obtaining a robust measurement for the entire volume, our analysis was performed on 50 sub-volumes of a uniform shape (128 × 128 × 128) randomly sampled from the input volume. The final effective spatial resolution reported for a given volume is the average of these individual SFSC measurements.

SFSC analysis was implemented using the open-source MIPLIB Python package (https://github.com/sakoho81/miplib, v1.0.6) for this study. The resolution cut-off was determined by a default criterion (SNR-0.5) recommended from official examples, and the obtained directional resolution profiles were visualized as a polar plot. For comparison among EM volumes, we used the resolution cut-offs on the 90° axis (corresponding to the XZ plane), on which this value was expected to be equally sensitive to image quality and slice thickness.

### Sensitivity analysis of effective resolution on SFSC threshold

To verify that changes in effective resolutions are robust to the choice of SFSC threshold, three commonly used cut-off criteria were applied to all synthetic FIB-SEM datasets in parallel with the default SNR-0.5: the empirically determined threshold (0.143) widely adopted in single-particle cryo-EM analysis, and the half-bit criterion derived from information theory. The effective XZ resolutions obtained under all three criteria scaled in the same manner with axial sampling rate and dwell time (Figure S2).

### Quantification of automatic mitochondrial segmentation

The data quality was assessed based on the mitochondrial segmentation accuracy of a published 3D AI segmentation model combining a U-Net with Swin-Transformer blocks (STU-Net)^39^. Models were trained independently for down-sampled image stacks of isotropic voxel sizes (5, 10, and 15 nm). The training data and corresponding ground truth were provided by the aforementioned FIB-SEM dataset. The training was conducted for 200 epochs using a Stochastic Gradient Descent (SGD) optimizer with momentum of 0.99 and a fixed learning rate of 1 × 10^-3^. The model from the final epoch was used for inference. Each model was applied to its corresponding set of testing data with the same pixel size, and the segmentation performance was quantified on a slice-by-slice basis by calculating the Intersection-over-Union (IoU) against the human ground truth, which was provided by the same public dataset and processed with the corresponding down-sampling factors. We plotted the segmentation error as 1-IoU (Figure 1E) to ease visual comparison with the effect resolution of the corresponding dataset (Figure 1D).

### AI-based Z-interpolation of serial images

The input image stack was acquired at a 2-μs dwell time and spaced at 100 nm in Z (Figure 2B). For the interpolation, we fed the pretrained Real-Time Intermediate Flow Estimation (RIFE) model^40^, which was previously shown to have outstanding performance on the FIB-SEM data,^41^ with the input stack. By a 2-fold interpolation, the RIFE artificially improved the Z-spacing of the original dataset from 100 nm to 50 nm.

### Quantification of inter-slice structural continuity

To quantitatively evaluate the 3D structural continuity, we calculated the mean Structural Similarity Index (SSIM)^42^ between consecutive slices, using the implementation provided by pyiqa (https://github.com/chaofengc/IQA-PyTorch<u>, v0.1.14.1</u>). The metric assesses the perceptual similarity between two images (*x* and *y*) by comparing luminance, contrast, and structure. This metric is defined as:

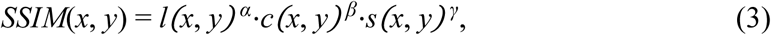

where luminance comparison 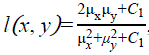, contrast comparison 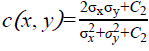, and structure comparison 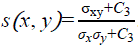. The terms 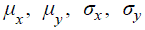 and *σ_xy_* are the local means, standard deviations, and cross-covariance for the corresponding images. The constants *C*_1_=(*K*_1_*L*)^2^, *C*_2_=(*K*_2_*L*)^2^, and *C*_3_=*C*_2_/2 stabilize the calculation, where *L* is the dynamic range of the pixel values. In our experiments, this metric was implemented using parameters *K*_1_=0.01 and *K*_2_=0.03. This calculation was applied in two distinct ways:

1. Independent analysis: To compare the overall continuity of different restoration methods, 50 sub-volumes (256 × 256 × 24) were independently and randomly sampled from each full volume. The mean SSIM was calculated between every two images within each cropped sub-volume to generate a distribution as a continuity measure.
2. Paired analysis: To compare 2D-denoising and v-denoising as a function of cutting thickness (25 nm to 200 nm), a paired-sampling analysis was conducted. For each thickness, 50 sub-volumes (256 × 256 × 24) were randomly sampled from identical coordinates in both the 2D-denoised and v-denoised image stacks. The mean SSIM was calculated for each sub-volume, resulting in 50 paired scores for statistical testing. The calibration baseline (dotted line) was established by sampling 500 random, non-adjacent slice pairs from volumes and computing the mean SSIM.

### Volumetric denoising network architecture and model training

For volume denoising, we adopted a cascaded U-Net-based deep denoising model named GShiftNet^43^ as the core architecture (Figure S3). The proposed model contained an initial feature extraction block enhanced by channel attention, followed by an encoder for fusing multi-frame features by channel-wise circular shifts and 8-direction spatial shifts. The processed features were then passed through three cascaded U-Net sub-networks for deep representation learning. Throughout the network, Channel Attention Blocks (CABs) refined features by adaptively reweighting channel responses via global average pooling, making the model focus on informative signals while maintaining stability through residual connections. The final reconstruction module aggregated multi-stage features and combined them with the input via a global residual path, followed by a convolutional layer to produce the denoised output.

All volumetric denoising training was performed using 3D patches of a fixed size (128 × 128 × 5). While the initial patches were all randomly sampled from the data volumes, the precise method of generating the training pairs differed for each workflow, as detailed in the following subsections. Standard data augmentation techniques, including random flips and rotations, were applied to each patch prior to the network feeding. The 540 training pairs and 32 validation pairs per epoch were sampled exclusively from the training region; the validation pairs functioned as a convergence-monitoring diagnostic and not as a hold-out for model selection. All models were trained for a fixed 100 epochs, and the model checkpoint from the final epoch was used for all subsequent inference. During training, the model was optimized using a weighted sum of the L1 and structural similarity loss (defined as 1–SSIM). The loss function *L_total_* used in training is defined as:

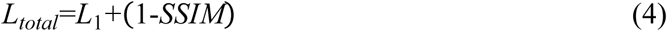

Training was performed with Adam (initial LR 2 × 10-4, β₁ = 0.9, β₂ = 0.99) and a plateau-based learning rate scheduler. All experiments were run on a Linux server equipped with a single Nvidia RTX 4090 GPU.

### Image preprocessing for volumetric denoising network training and inference

All raw 8-bit VEM images were globally normalized to the [0, 1] range by dividing pixel intensities by 255 prior to both training and inference. No min-max rescaling, percentile clipping, or per-image standardization was applied, so that the relative intensity distribution of the raw images was preserved across slices and across datasets. The same normalization was applied identically to inputs, supervision targets (when applicable), and inference inputs. After inference, denoised outputs in the [0, 1] range were rescaled back to 8-bit by multiplying by 255 for downstream visualization and analysis.

### Supervised training of the volumetric denoising model

For the supervised denoising experiments (Figure 3), 61 full-frame slices (4000 × 4000 pixels each) of the reference, snapshot, and hybrid stacks were used for training. Two distinct data preparation pipelines were implemented to generate source and target pairs to train our denoising model:

1. For the fast and slow imaging of S/r-stack, whose input sequence consisted of five consecutive frames (the 1^st^ and 5^th^ frames were captured by slow-scan; the 2^nd^ to 4^th^ frames were from fast scan), an approach was employed to prevent the model from learning a trivial identity map for the slow-scanned frames. To construct paired training samples, a larger patch of size (256 × 256 × 5) was initially extracted from both the hybrid stack and the corresponding high-SNR ground truth. From these, the input and supervision patches (128 × 128 × 5) were derived using a deterministic checkerboard subsampling scheme: the input patch was composed of pixels located at even rows and even columns (top-left), while the supervision patch was constructed from the diagonally opposite subset (bottom-right). This design enabled robust end-to-end denoising of hybrid patches.
2. For the fast-scan mode, training pairs were generated by directly extracting a 3D patch (128 × 128 × 5) from the snapshot stack and pairing it with the corresponding patch from the ground truth at the same spatial location. This pairing strategy enabled the model to learn a direct mapping from noisy inputs to their clean counterparts. For the comparable 2D supervised denoising model (Fig. 3), the training process was identical, except that the input patch depth was reduced to one (128 × 128 × 1). This minimal modification ensured a fair comparison between the 2D and 3D approaches by keeping the model architecture, number of training samples, and optimization strategy consistent.

### Self-supervised training of the volumetric denoising model

For the self-supervised denoising experiments lacking paired low-noise references (Figure 4), training pairs were generated directly from the raw images. For each thickness, 61 full-frame slices separate from the snapshot dataset were used as a training source. The pairing strategy followed the same principle as the supervised model training scheme described above. Specifically, a randomly cropped patch (256 × 256 × 5) was divided into two sub-patches using a deterministic checkerboard pattern, which were then assigned as the source and target, respectively. This blind-spot design encouraged the network to learn underlying structural representations rather than simply fitting noise patterns.

### Model inference of denoised volumetric datasets

For inference on large volumes, a patch-based approach was implemented using an overlapping sliding window strategy. To prevent block artifacts in the denoised stacks, adjacent patches (128 × 128 × 5) were processed with a 12.5% overlap in the XY plane (16 × 16), and 20% (one layer) overlap in the Z-axis. The final denoised pixels in these overlapping regions were the mean of the denoised outcomes from the adjacent patches. While this inference pipeline was applied to all denoising strategies, the hybrid fast-slow mode included an additional post-processing step to ensure a fair comparison with the interpolation method. After the initial seamless volume was generated, the slices corresponding to the positions of the original slow-scan frames (e.g., the 1^st^ and 5^th^ for every patch) were replaced with the original, unprocessed high-SNR slow-scan images. This step guaranteed that both volumetric denoising methods and the RIFE interpolation method share identical data at the ground truth layers.

### Mitochondrion and synapse segmentation of authentic datasets

In Figure 5, mitochondria and synapses were manually annotated in the reference dataset (2.0-µs dwell time) by an expert annotator. These annotations serve either as training data for the segmentation models or ground truth for evaluating the segmentation accuracy. To ensure that the test data were entirely unseen by the models, training and test datasets were chosen from separate subregions of the volume.

Next, the raw dataset acquired at 0.5 µs/pixel was denoised using four different methods: 3 × 3 mean filter of Fiji, 2D self-supervised Blind2Unblind, self-supervised DeepSeMi, and volumetric denoising (v-denoising). All three deep learning-based methods (Blind2Unblind, DeepSeMi, and v-denoising) were trained on the same volume using the reference data.

For the comparison of segmentation accuracy, the same segmentation pipelines were applied to the denoised outputs as well as the raw and reference datasets. Specifically, automatic segmentation of mitochondria and synapses was performed in ZEISS arivis Cloud, a web-based AI segmentation platform (https://www.zeiss.com/microscopy/en/products/software/arivis-cloud.html). For each target structure (mitochondria and synapses), a custom deep-learning segmentation model was trained on the platform using its default pipeline; the model architecture, training schedule, and prediction process were managed automatically by the platform. Both segmentation models were trained on the reference volume of a different region than the test volume and were manually annotated by an expert. Each trained model was then applied without further re-training to all test volumes (the snapshot, four denoised outputs, and the reference), all corresponding to the same spatial test region. Segmentation outputs were visualized in ZEISS arivis Pro v4.4.0 (https://www.zeiss.com/microscopy/en/products/software/arivis-pro.html) and quantitatively compared with the manual ground truth on the test region by computing Intersection-over-Union (IoU) slice by slice.

### Automated neurite segmentation

A sub-volume of 1000 × 1000 × 1000 voxels (12 × 12 × 35 nm) was extracted from a previously published SBF-SEM dataset of adult zebrafish spinal cord and processed in parallel as the raw stack and as a volumetric denoised stack (using the self-supervised training pipeline described above). Automated neurite reconstruction was performed on both datasets using an identical model. The volume was first partitioned along the Z-axis into non-overlapping sub-volumes of 200 consecutive sections to enable scalable block-wise processing. Each sub-volume was then independently processed by the SegNeuron^44^ generalist model to predict voxel affinities between neighboring locations. Based on the affinity maps, seeded watershed was applied to generate an initial over-segmentation into supervoxels, which was subsequently refined through multicut-based agglomeration^45^ to recover topologically coherent neuronal fragments within each block. Finally, segmented instances from adjacent sub-volumes were reconciled and stitched using an overlap-based matching strategy, yielding globally consistent neuron reconstructions across the full sub-volume.

### Manual proofreading and time measurement

Seven neurites were selected randomly from each of the two reconstructed volumes for manual proofreading, and the same set of seven neurites (matched by location) was traced in both the raw and denoised reconstructions. Proofreading and visualization were performed in ZEISS arivis Pro v4.4.0 by a single expert annotator. To control for potential learning, fatigue, and order effects, the proofreading order of raw and denoised reconstructions was counterbalanced across the seven neurites. Wall-clock time was used to measure the duration of each proofreading session, and over-segmentation counts and proofreading times were normalized to the path length of each neurite for cross-neurite comparison.

## QUANTIFICATION AND STATISTICAL ANALYSIS

All statistical tests were performed using GraphPad Prism 10.1.2 built-in functions. We used two-tailed paired t-tests (Figure 3D), two-sample t-tests (Figures 3F, 4D), two-tailed unpaired t-tests (Figure 4D and 4G), Friedman tests with Dunn’s post-hoc multiple-comparison correction (Figures 3E, Figure 5C and 5E), and paired Wilcoxon signed-rank tests (Figure 6C and 6D). All Data are presented as the mean ± s.d., with statistical significance (*p < 0.05, **p < 0.01, ***p < 0.001, ****p < 0.0001).

### Supplemental video and Excel table titles and legends

**Supplemental video 1. Imaging dwell time-dependent cutting artifacts**

**Supplemental video 2. Volumetric denoised output – XY view**

**Supplemental video 3. Volumetric denoised output – XZ view**

